# BCR-ABL promotes hematopoietic stem and progenitor cell formation in embryonic stem cells

**DOI:** 10.1101/2023.02.03.526926

**Authors:** Jérôme Artus, Alina Zenych, Isidora Simanic, Christophe Desterke, Denis Clay, Sonia Saïm, Yousef Ijjeh, Lucas Eduardo Botelho de Souza, Sabrina Coignard, Annelise Bennaceur-Griscelli, Ali G. Turhan, Adlen Foudi

## Abstract

Generating Hematopoietic Stem Cells (HSCs) from Pluripotent Stem Cells (PSCs) has been a long-lasting quest in the field of hematopoiesis. Previous studies suggested that enforced expression of BCR-ABL, the unique oncogenic driver of Chronic Myelogeneous Leukemia (CML), in Embryonic Stem Cells (ESCs)-derived hematopoietic cells is sufficient to confer long-term *in vivo* repopulating potential. To precisely uncover the molecular events regulated by the Tyrosine-kinase activity of BCR-ABL1 (p210) during the course of hematopoietic differentiation, we engineered a Tet-ON inducible system to modulate its expression in murine ESC. We showed in unique site-directed knock-in ESC model, that *BCR-ABL* expression tightly regulated by doxycycline (dox) controls the formation and the maintenance of immature hematopoietic progenitors. Interestingly, these progenitors can be expanded *in vitro* for several passages in the presence of dox. Our analysis of cell surface markers and transcriptome compared to wild-type fetal and adult HSCs unraveled a similar molecular signature. LTC-IC assay confirmed their self-renewal capacities albeit with a differentiation bias towards erythroid and myeloid cells. Collectively, our novel Tet-ON system represents a unique *in vitro* model to shed lights on ESC-derived hematopoiesis, CML initiation and maintenance.

**KEY POINTS:**

- We report a unique BCR-ABL-induced-embryonic stem cell -derived hematopoiesis model in murine embryonic stem cells
- This BCR-ABL-induced self-renewal and differentiation model can be of major interest to uncover molecular events required for ESC-derived hematopoiesis

## INTRODUCTION

Hematopoietic Stem Cells (HSCs) ensure the continuous formation of blood cells throughout the life of an individual due to their unique properties of self-renewal and differentiation. Pluripotent Stem Cells (PSCs), including Embryonic Stem Cells (ESCs) and induced pluripotent stem cells (iPSCs), are characterized by their ability to generate all embryonic cell lineages. Generating HSCs from PSCs would provide a powerful paradigm to uncover molecular mechanisms regulating normal hematopoiesis, to model hematological disorders and engineer cell-based therapies. However, the identification of true long-term HSCs with both self-renewal and multipotent differentiation abilities from PSCs remains a major challenge. This limitation could be partly due to our limited knowledge of the ontogeny of HSCs, the lack of defined permissive conditions to drive PSC differentiation towards an HSC phenotype and the inability to maintain and expand them. Among the numerous attempts, distinct but not exclusive strategies have been commonly used including recapitulating embryonic hematopoiesis in a dish, direct conversion by enforced expression of hematopoiesis-lineage specific regulators, teratoma formation or direct cell lineage conversion (for review see ^1^).

Chronic Myeloid Leukemia is a myeloproliferative disease characterized by an excessive proliferation and differentiation of myeloid cells and a long-term clonal maintenance of leukemic stem cells (LSCs) driven by BCR-ABL expression in HSCs ^2^. LSCs can self-renew and exhibit multipotent lymphoid and myeloid potential with altered bias towards myeloid differentiation during the chronic phase. Retroviral transduction of BCR-ABL into hematopoietic bone marrow cells confers CML-like disease when transplanted into mice. Importantly, transduced cells are able to reconstitute both lymphoid and myeloid cell lineages ^3^. Similar transduction experiments were performed in mESCs and mESC-derived hematopoietic cell lines were established ^4^. They represent primitive hematopoietic progenitors with *in vitro* and *in vivo* leukemic potential and limited differentiation capacities. Interestingly, mESCs over-expressing BCR-ABL retains multipotent hematopoietic differentiation potential with a long-term repopulation potential in a transplantation assay ^5,6^. BCR-ABL expression during ESC differentiation alters the proliferation and differentiation of immature progenitors in vitro ^5,7^.

To get a better understanding of the role of BCR-ABL during the course of ESC differentiation, we generated a unique Tet-On inducible system to regulate BCR-ABL expression in a Doxycycline-inducible manner. As compared to other ESC-derived models, the model that we describe here is characterized by a site-directed knock-in strategy at the ESC stage, allowing to analyze the effects of BCR-ABL expression at various stages of hematopoiesis formation. We observed that BCR-ABL expression favors the generation and the maintenance of immature hematopoietic progenitors. Interestingly, these progenitors can be maintained *in vitro* for an extended period of time if BCR-ABL expression is sustained with cellular dynamics affected by IL-6 signaling. Cell surface marker analysis and transcriptome comparison with fetal and adult LSK cells unravel a similar molecular signature to hematopoietic stem and progenitor cells. In addition, we tested their self-renewal capacities with LTC-IC assay and observed biased differentiation abilities towards erythroid and myeloid cell lineages at the expense of lymphoid cell lineages. Overall, the model that we report represents a unique *in vitro* cellular tool to shed light on ESC-derived hematopoiesis, leukemia initiation and leukemic stem cell maintenance.

## METHODS

### Generation of *BCR-ABL^P210^* inducible ESCs

*BCR-ABL^P210^* coding sequence was recovered from *Eco*RI digest of pMIGR-BCR-ABL-IRES-GFP plasmid ^8^ and ligated into *Eco*RI-linearized pBS31 flip-in vector ^9^.

KH2 ESC ^9^ were electroporated with pBS31-BCR-ABL^p210^ and FLP recombinase plasmids (NEON transfection system, Thermo Fisher) and selected with Hygromycin (Invitrogen). After 12 days, hygromycin-resistant ESC colonies were manually picked, expanded and genotyped by PCR (HotStar, Qiagen) to confirm the presence of *BCR-ABL^p210^* coding sequence and the genetically modified alleles at the *ColA1* and *ROSA26* loci. We used primers sequences available at the Jackson Laboratory to genotype *Cola1* (18465, 18466 and IMR6726) and *ROSA26* (IMR8052, IMR8545 and IMR8546). *BRC-ABL* primers were: ABLK-F: 5’-CGCAACAAGCCCACTGTCT, and ABLK-R: 5’-TCCACTTCGTCTGAGATACTGGATT. 13 out of 13 ESC clones were analyzed and all possessed the correct genotypes.

### Cell culture

Murine ESC (KH2) and EBs were cultured as described in ^10^. To establish HSC Lines, day 5-EBs were dissociated with 0.25% Trypsin-EDTA (Gibco) and plated at ~2×10^4^ cells/cm^2^ on mitomycin C-treated OP9 cells (ATCC) in HSCL medium containing Minimal Essential Medium Alpha medium supplemented with 20% FBS, 2 mM L-Glutamine, 100 units/mL Penicillin/Streptomycin (Gibco) and 1 μg/mL dox. Cells were passaged onto new OP9 cells 10-14 days later. Once established, HSCL were routinely passaged using PBS-EDTA dissociation solution (0.5 mM-EDTA, 0.9 g/L NaCl) every 2-3 days. They were seeded at ~2×10^4^ cells/cm^2^ on OP9 cells in HSCL medium.

For lymphoid differentiation, HSCL or fetal liver LSK cells were cultured for 2 weeks on OP9 or OP9-DL4 cell layer in HSCL medium supplemented with 10 ng/mL Flt-3 (PeproTech) and 10 ng/mL IL-7 (PeproTech).

### Colony formation cell assay

CFC assays were performed using ES-Cult (M3120, StemCell Technologies) for EB-derived cells and Metho-Cult (M3234, StemCell Technologies) supplemented with cytokines for HSCL according to the manufacturer. Colonies were scored 12-14 days later.

### Flow cytometry

Single cell suspensions were pre-incubated for 15 minutes in a staining solution (PBS+/+, 2% FBS) containing 1 μg/mL Fc block (BD Biosciences), followed by an incubation with the antibodies for 45 minutes (Table S1). To discriminate live from dead cells, 1 μg/mL propidium iodide (Invitrogen) was used. Flow cytometry analysis were conducted using BD LSR Fortessa™ cell analyzer or MACSQuant^®^ analyzer (Miltenyi Biotec). Data were prepared using FlowJo 10.1 (Tree Star, Inc.) software.

### LTC-IC

Single cell LTC-IC assays were performed according to ^11^. Single ESLAM cells were sorted using BD FACSAriaTM II cell sorter (BD Biosciences) and co-cultured on AFT024 feeder during 4 weeks, in LTC-IC culture condition, before CFC assessment in methyl-cellulose.

### Microarray analysis

LSK cells from adult bone marrow, E14.5 fetal liver and HSCL#A and #C were sorted in triplicates. Total RNAs were extracted using PureLink^®^ RNA Mini Kit (Thermo Fisher Scientific) and treated with DNase using On-column PureLink^®^ DNase Treatment (Thermo Fisher Scientific). The quality of the samples was determined using a Bioanalyzer 2100 (Agilent Technologies). cDNAs were obtained using Ovation^®^ Pico WTA System V2 (NuGEN Technologies) and hybridized on GeneChip™ Mouse Gene 2.0 ST Array (Affymetrix™). Microarray transcriptome experiments were normalized with (Robust Multi-Array Average (RMA) algorithm ^12^. Data can be accessed at Gene Expression Omnibus accession GSE198018.

Bioinformatics analyses were performed in R software environment version 4.1.0. Unsupervised selection of most variable genes was performed with custom R function: “varsig” available at the following address: https://github.com/cdesterke/matrixtools (accessed on 2022, january 6th). This function takes as input parameter the expression matrix and returns the most variable genes between samples with a variance over the global variance of the array. Principal component analysis on variable genes was performed with FactoRineR R-package version 2.4. Expression heatmap was drawn with pheatmap R-package version 1.0.12 using Euclidean distances and ward. D2 method. Expression boxplot were drawn with ggplot2 version 3.3.5. Gene set enrichment analysis was done with GSEA standalone software version 4.0.3 ^13^ on MsigDb database version 7.0 ^14^.

## RESULTS

### BCR-ABL^p210^ expression promotes the formation of immature hematopoietic progenitors

To address the possibility that *BCR-ABL^p210^* expression might affect ESC-derived hematopoiesis, we generated a genetically modified mESC harboring a tetracycline inducible system allowing the tight control of *BCR-ABL^p210^* expression (Fig. S1A). We first validated the functionality of the system by western-blot analysis (Fig. S1B). While in the absence of doxycycline p-BCR-ABL was not detected, 24h culture in presence of doxycycline was sufficient to induce the expression of p-BCR-ABL as well as an increase of p-CRKL, one of the surrogate targets of BCR-ABL tyrosine kinase activity.

We next cultured ESC without LIF under non-adherent conditions where they form Embryoid Bodies (EBs), which recapitulate several developmental events including hematopoiesis ^15,16^ (Fig. 1A). Doxycycline was added at several time points starting at day 0, day 5 (which corresponds to the time when hematopoietic precursors start to be formed ^17,18^ or at day 10 of EB differentiation. After 10 days, the number of EB-derived hematopoietic precursors was determined in CFC assays (Fig. 1A, B). We first observed a significant 2-fold reduction of the number of CFC colonies when the *BCR-ABL^p210^* expression is induced from day 0 in ESC. In sharp contrast, when doxycycline was added from day 5, the number of hematopoietic colonies increased by 3.7-fold. Our observation suggested that an early expression of *BCR-ABL^p210^* expression in mESCs at pluripotent stage have a detrimental effect on hematopoietic differentiation. We thus added doxycycline during the 5 first days of differentiation and observed a moderate, although non-significant, reduction of the number of hematopoietic colonies by 1.5-fold. To dissect more precisely the time window when BCR-ABL^p210^ exerts its beneficial effect, we treated EBs from day 5 to day 10 or starting from day 10 with doxycycline. We observed in both conditions, an increase of hematopoietic colonies by 1.8-fold. Altogether, our results confirm that *BCR-ABL^p210^* expression from day 5 promotes the commitment of EBs into hematopoietic progenitors.

**Figure 1.**
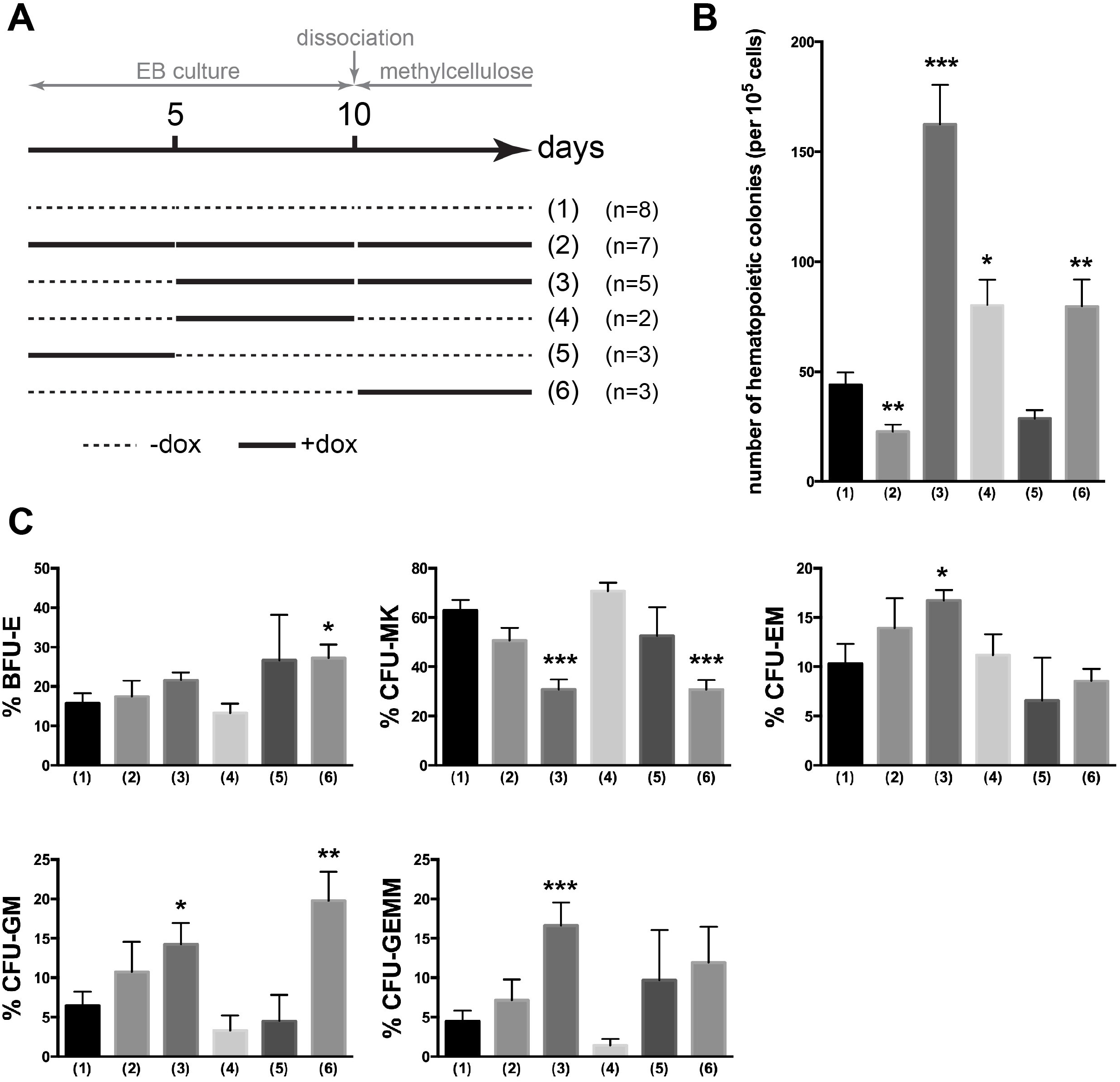
*BCR-ABL1^p210^* expression affects the formation of hematopoietic progenitors from ES cells. **(A)** Schematic representation of ESC differentiation and CFC assay. **(B)** Number of hematopoietic colonies after 12-14 days in methylcellulose. **(C)** Number of CFU-E, CFU-MK, CFU-EM, CFU-GM and CFU-GEMM colonies after 12-14 days in methylcellulose. n: number of experiments. Statistical Mann–Whitney tests are indicated when significant (*, p < 0.05; **, p < 0.01; ***, p < 0.001).

Subsequently, we determined the differentiation potential of hematopoietic progenitors consecutive to *BCR-ABL^p210^* expression, in CFU colonies assays (Fig. 1C). Among the noticeable differences, we showed that EBs treated from day 5 with doxycycline gave rise to an increased proportion of CFU-EM (erythroid/megakaryocyte), CFU-GM (granulocyte/macrophage) and CFU-GEMM (granulocyte/erythroid/macrophage/megakaryocyte) colonies and reduced proportion of CFU-Mk (megakaryocyte), as compared to untreated conditions. EBs treated from 10 days produced less CFU-Mk and more BFU-E and CFU-GM colonies. Taken together, we conclude that the timing of *BCR-ABL^p210^* expression during the course of EB differentiation enhance the number and the differentiation potential of hematopoietic progenitors. Particularly, *BCR-ABL^p210^* expression at day 5 favors the emergence of immature hematopoietic progenitors harboring multipotent differentiation potential (CFU GEMM).

### BCR-ABL^p210^ expression allows the expansion of hematopoietic cell lines

We hypothesized that *BCR-ABL^p210^* expression starting from day 5 might also favor the proliferation of hematopoietic progenitors in permissive culture conditions. EBs were dissociated at day 5 and plated on OP9 stromal cells in presence of doxycycline (Fig. 2A) without any added cytokines. In this culture condition, we observed an expansion of small clusters of semi-adherent cells, organized as blastic-like immature colonies (Fig. 2B). Immature and undifferentiated blast cells harboring few signs of differentiation were confirmed by morphology after MGG staining (Fig. 2C). Interestingly, these progenitors exhibited sustained proliferation for extended periods of time (Fig. 2D), from which two cell lines were established (hereafter referred as HSCL#A and HSCL#C).

**Figure 2.**
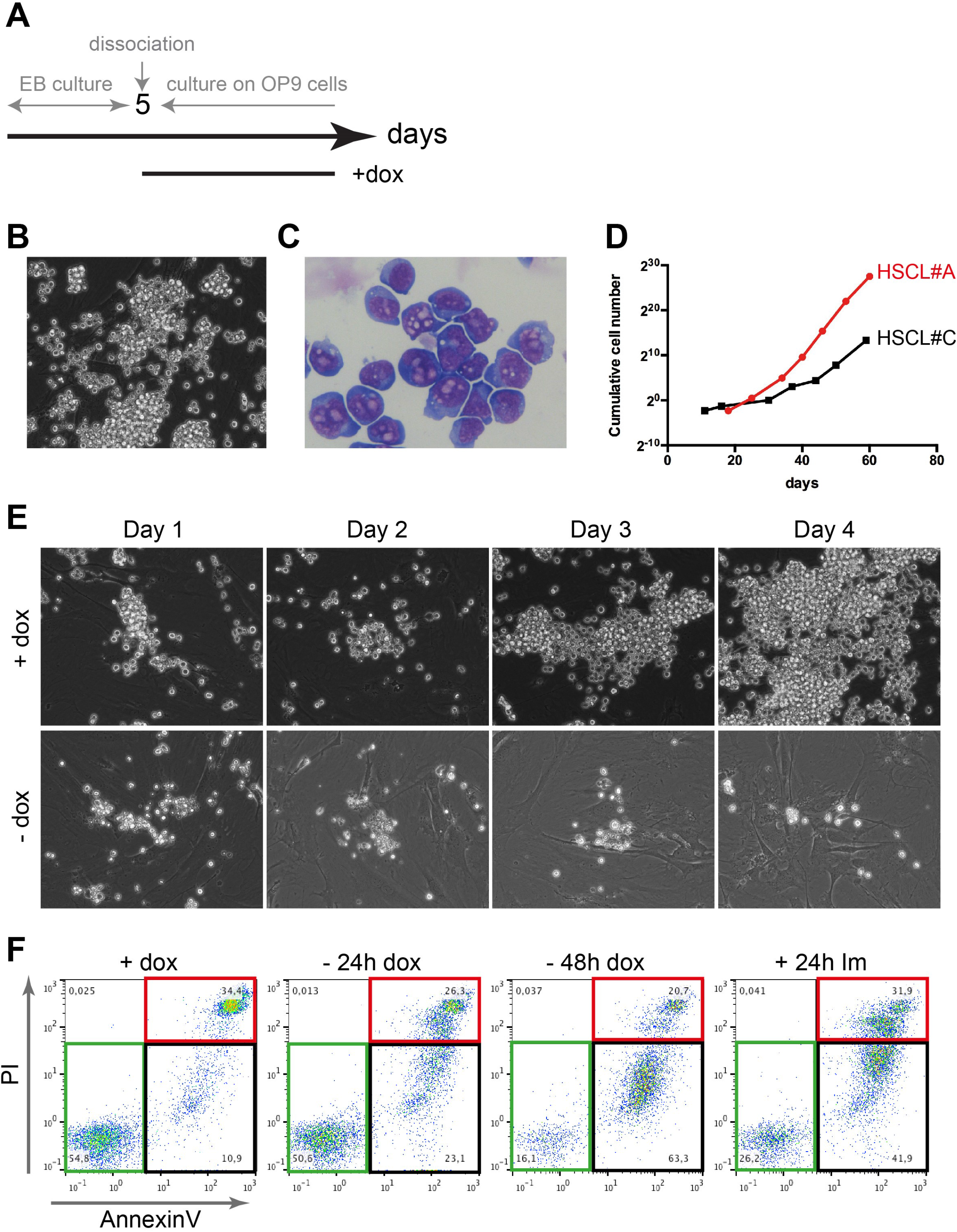
ES cell-derived hematopoietic cell lines can be established under *BCR-ABL^p210^* expression. **(A)** Diagramatic representation of the timeline of ESC differentiation protocol. **(B)** HSCL#A cell morphology. **(C)** MGG staining of HSCL#A cells. **(D)** Proliferation curves of HSCL#A and HSCL#C cell lines over 40 days in culture. **(E)** Kinetics of the effect of doxycycline removal on HSCL#A cells cultured on OP9 cells for 4 consecutive days. **(F)** Flow cytometry of Annexin V and PI stained HSCL#A cells. Cells were cultured in presence of doxycycline (+dox), 24h (-24h dox) and 48h (-48h dox) after doxycycline removal or in presence of 10μM Imatinib for 24h (+24h Im). (B, E) Images were acquired using a 20X objective.

As BCR-ABL oncogene induces the survival of leukemic cells ^19,20^ in CML, we tested whether HSCL cell growth is related to *BCR-ABL^p210^* expression. We analyzed the cell behavior following doxycycline withdrawal for 4 consecutive days (Fig. 2E). While HSCL#A cells grew rapidly in presence of doxycycline, HSCL#A failed to proliferate and died in its absence, confirming dependence to BCR-ABL activity and their non-autonomous proliferation and/or survival capacity. We analyzed by flow cytometry the AnnexinV-positive/PI-negative cell population corresponding to early apoptotic events (Fig. 2F). We observed an increase of the AnnexinV-positive/PI-negative cell population 24h after doxycycline removal (23.1% without *vs*. 10.9% with dox). This effect was even more pronounced 48h after (63.3%). In addition, we also observed apoptosis events in conditions where HSCL#A cells were cultured in presence of doxycycline and the Tyrosine Kinase inhibitor Imatinib, during 24h (Fig. 2F).

Taken together, TK-activity of BCR-ABL oncogene favors the emergence, selfrenewal and long-term maintenance/survival of immature hematopoietic progenitors derived from mESCs.

### HSCL cell lines exhibit a molecular signature resembling HSPC

We further characterized the phenotype of these HSCLs (Fig. 3 and Fig. S2). HSCs reside in the Lin-, Sca-1+, c-Kit+ population (LSK). We show that HSCL#A cells are c-Kit+ (CD117) and 10-20% Sca-1+ and lack the expression of hematopoietic lineage (Lin-) markers, such as TER-119 (erythroid cells), Mac-1 (monocytes/macrophages), Gr-1 (granulocytes), B220 (B lymphocytes), and CD3e (T lymphocytes) (Fig. 3A). In addition, we analyzed the cell surface markers of the SLAM family receptors that delineate HSCs from MPPs ^21^. Interestingly, most of the HSCL#A cells were CD150+, CD48- (81%) independently from their expression of Sca-1 (81% CD150+ CD48- LSK and 95% CD150+ CD48- LK). HSCL#C cells had a slightly different distribution with 90% of LSK and 57% of CD150+ CD48- LSK and 70% of CD150+ CD48- LK (Fig S2A). We also used another panel of surface markers (ESLAM: EPCR, CD45, CD48) that delineate HSCs ^22^. Compared to mHSCs in BM, EBs-derived HSCL#A cells expressed lower levels of CD45 and higher levels of EPCR (Fig. 3B). HSCL#C cells had a similar distribution CD48- CD45Lo EPCRHi (Fig. S2B). Lastly, we also demonstrated that they express low levels of CD34 and do not express CD135 (data not shown). Overall, these results show similar phenotype of HSCL with BM HSCs and immature multipotent progenitors.

**Figure 3.**
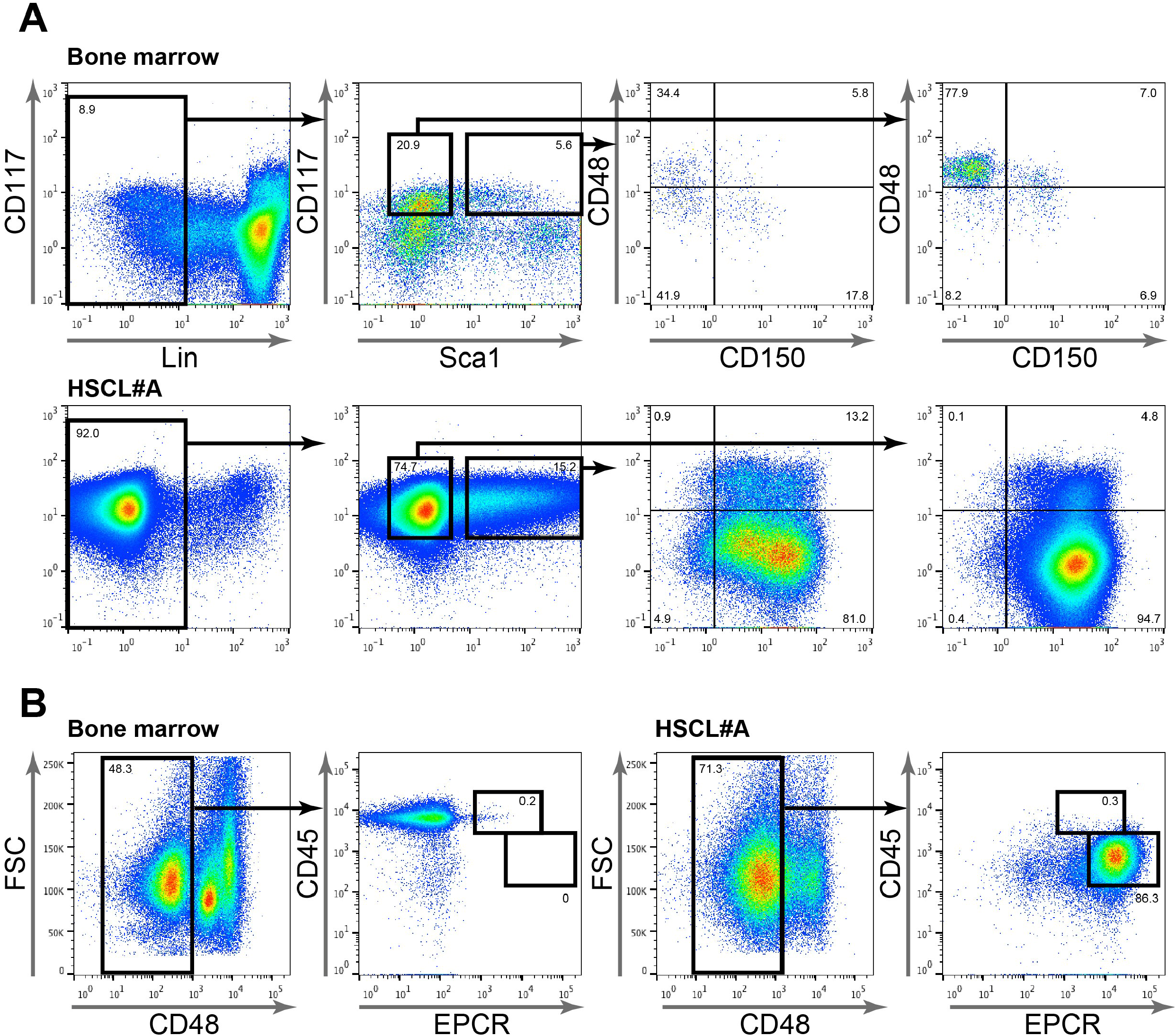
HSCL#A cells express surface markers characteristic of immature hematopoietic progenitors. **(A)** Representative FACS profile of CD48-CD150+ fraction in LSK (Lin-Sca1+CD117+) and LK (Lin-Sca1-CD117+) cell populations. **(B)** Representative FACS profile of CD48-CD45+EPCR+ populations. The gating strategies were designed according to the bone marrow samples.

The ontogeny of HSCs *in vivo* is a highly dynamic process, that occurs at different times and locations during the embryonic and fetal development ^23,24^. The generation of *in vitro* HSCs from ESCs with a multipotent and long-term engraftment potential has been quite challenging ^25–27^. We characterized and compared the gene expression profiles of HSCL with fetal and adult HSPCs. To do this, we sorted LSK cell population from HSCL#A, HSCL#C, E14.5 fetal liver and adult bone marrow and analyze their transcriptome signatures by RNA-Seq (Fig. 4A). Unsupervised principal component analysis with the 1333 genes exhibiting the highest variability reveals that HSCL#A and #C clustered together and exhibited distinct molecular signature from fetal and adult LSK (Fig. 4B).

**Figure 4.**
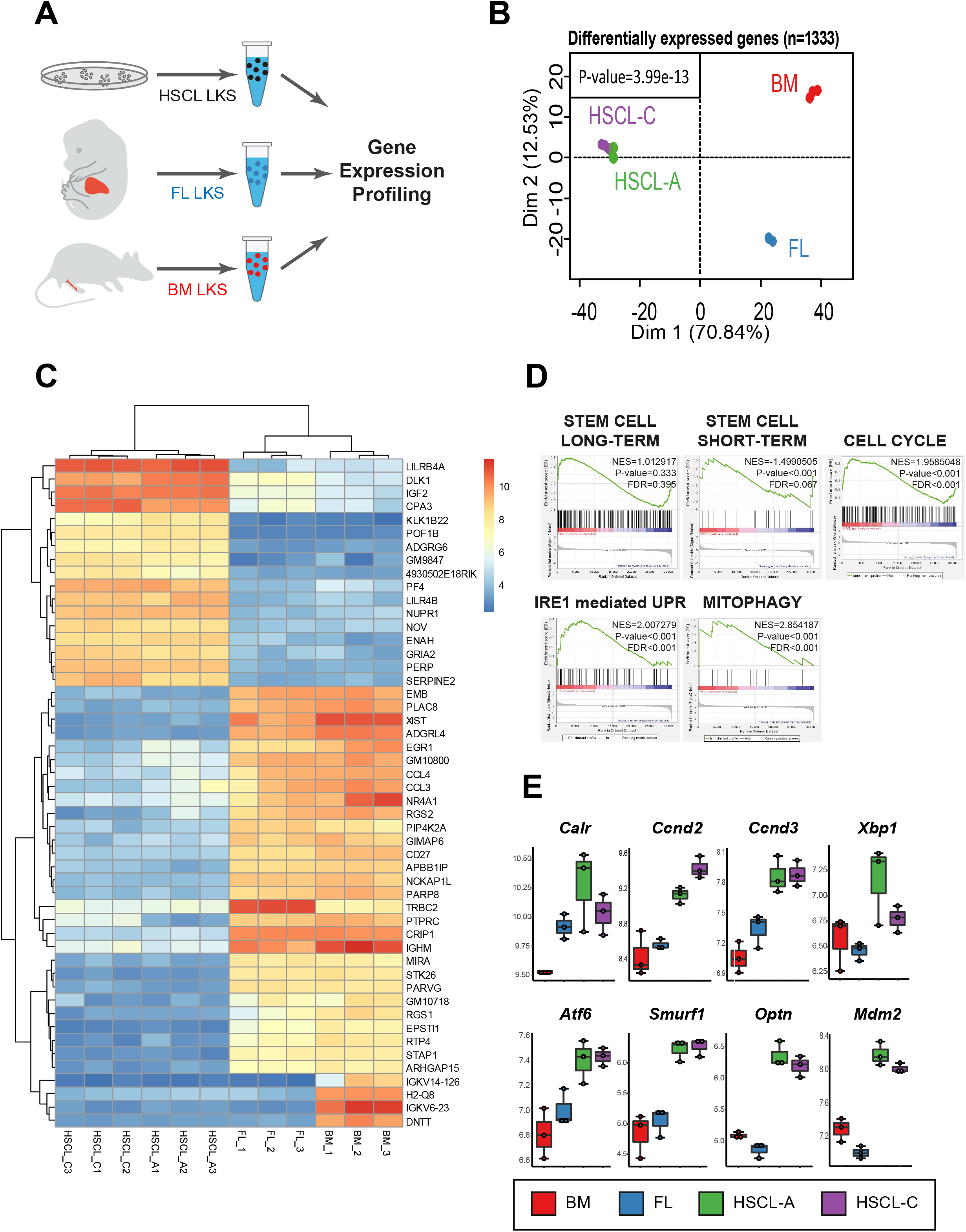
Gene expression comparison between LSK cells isolated from HSCL cell lines, fetal liver and adult bone marrow. **(A)** Schematic representation of the experimental design. **(B)** Principal component analysis (PCA) plot showing the distribution of the different samples according to the 1133 most variable genes among all the 12 microarray samples (p-value of group stratification on first principal axis). **(C)** Unsupervised classification and heatmap showing the expression of the most 50 variable genes. **(D)** Gene set enrichment analysis (GSEA) performed between HSCL samples versus other LSK samples with long-term HSC, short-term HSC, cell cycle, mitophagy gene sets. NES: normalized enrichment score, FDR: False Discovery Rate. **(E)** Expression boxplot of reliable genes identified during gene set enrichment analysis (log2 microarray expression). (B, E) purple: HSCL#C; green: HSCL#A; blue: FL; red: BM.

This stratification was validated by clustering with the 50 best variable genes (Fig. 4C). Among them, we found 17 genes highly expressed in HSCL which are associated to HSC quiescence (*Nupr1* ^28^, *Pf4* ^29^), myeloid differentiation (*Lilbr4a, Lilbr4b* ^30^) and LSCs (*Dlk1* ^31^, *Igf2* ^32^, *Pf4* ^33^). Several genes exhibited a lower expression in HSCL compared to adult and fetal LSK including genes associated to leukemia and LSC (*CD27* ^34^, *Crip1* ^35^, *Pip4k2a* ^36^, *Ptprc* ^37^, *Rgs1* ^38^, *Rgs2* ^39^, *Stap1* ^40^), HSC aging (*Egr1*, ^41^), myeloid differentiation (*Ccl3* ^42^, *Nckap1l* ^43^, *Rgs2* ^39^), lymphoid development (*Apbb1ip* ^44^, *Gimap6* ^45,46^, *Ighm* (IgM), *Nckap1l* ^45 43^*, Ptprc* (B220), *Trbc2*). Additionally, we found several lymphoid markers downregulated in both HSCL and FL LSK compared to BM LSK including the immunoglobulins IgKv14-126 and IgKv6-23, the Histocompatibility 2 Q region locus 8 (*H2-Q8*) and the deoxynucleotidyl transferase terminal (*Dntt*) ^47^. These observations could suggest an alteration of both myeloid and lymphoid differentiation programs and/or simply reflect the absence of lymphoid progenitors in the LSK cell population in HSCL.

Gene set enrichment analysis (GSEA) showed no significant difference in longterm HSC signature (NES=1.012917, p-value=0.333) and a decrease in short-term HSC signature (NES=-1.4990505, p-value<0.001) in HSCL (Fig. 4D). However, they exhibited a significant enrichment of cell cycle (NES=1.9585048, p-value<0.001, including *Ccnd2* and *Ccnd3*), unfolded protein response (NES=2.007279, p-value<0.001, including *Xbp1*) and mito/macro autophagy signature (NES=2.854187, p-value<0.001, including *Atf6, Smurf1, Optn*) (Fig. 4D and E). These observations support the idea that HSCL correspond to immature hematopoietic cells and unravel the activation of several stress pathways which have been shown to be misregulated in CML/AML LSCs, such as autophagy ^48^, mitophagy ^49^ and UPR ^50,51^.

### *In vitro* dynamic cell heterogeneity within HSCL cell population is driven by IL-6 cytokine signaling

Having established that HSCLs contain in culture both LK and LSK subpopulations, we next addressed whether culture conditions could modify this cell population heterogeneity. We first compared the LK/LSK distribution in HSCL#A cells cultured on OP9 stromal cells in presence or absence of five standard hematopoietic cytokines (SCF, IL-3, IL-6, EPO, TPO; 5GF) for 7 days (Fig. 5A). These five cytokines could promote the expansion of the LSK compartment (38.6% in presence of cytokines *versus* 22% in absence). We next determined which cytokine is important for the LSK compartment. We cultured HSCL#A cells without OP9 cells in presence or absence of a single cytokine (Fig. 5B). We observed a decrease of LSK cells in absence of cytokine and OP9 cells (8.3% in absence *versus* 22.0% in presence). The addition of single cytokine, SCF, IL-3, EPO or TPO did not promote the LSK compartment (7.3% with SCF, 6.7% with IL-3, 6.5% with EPO and 8.6% with TPO). In contrast, the addition of IL-6 strongly increased the LSK compartment (34.3%). Thus, IL-6 seems to impact the equilibrium between LK and LSK cell populations.

**Figure 5.**
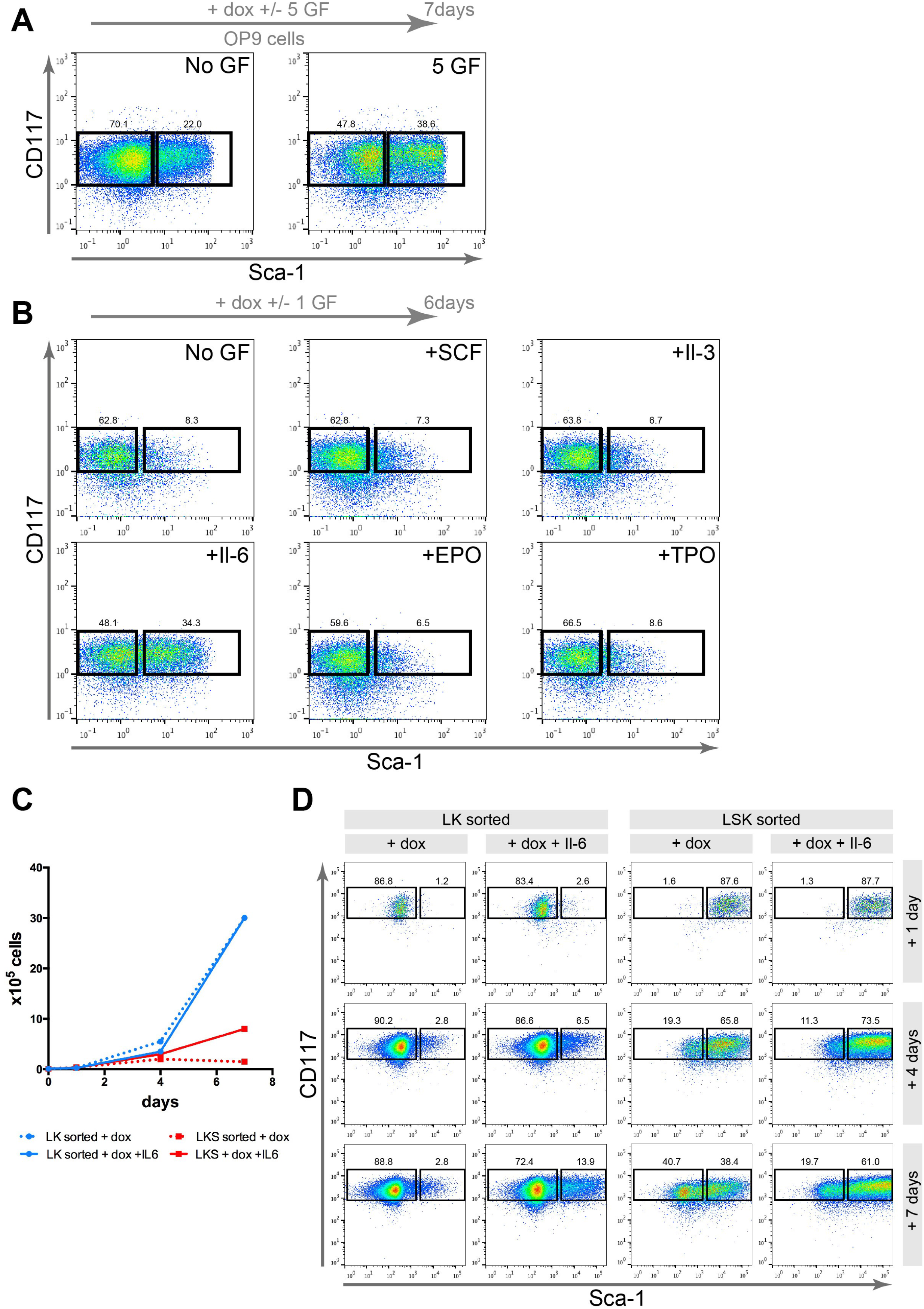
Il-6 promotes the expansion of LSK compartment in HSCL#A cell line. **(A)** FACS profile of CD117 and Sca-1 in Lin- cell populations of HSCL cultured on OP9 and doxycycline in presence or absence of SCF, Il-3, Il-6, EPO and TPO (5 GF) for 7 days. **(B)** FACS profile of CD117 and Sca1 in Lin- cell populations of HSCL cultured with doxycycline in presence or absence of SCF, Il-3, Il-6, EPO or TPO for 6 days. **(C)** Proliferation curves over a 7 days culture period of LK and LSK cell sorted cell populations with doxycycline in presence or absence of Il-6. **(D)** FACS profile of CD117 and Sca-1 in Lin- cell populations of LK and LSK cell sorted populations cultured with doxycycline in presence or absence of Il-6 after 1, 4 and 7 days in culture.

We next tested whether IL-6 signaling might regulate differentially LK and LSK cells. To test this hypothesis, we sorted LK and LSK cells and cultured them in presence or absence of IL-6 for 7 days. On days 1, 4 and 7, we counted the number of cells (Fig. 5C) and analyzed by flow cytometry the distribution of LK and LSK respectively (Fig. 5D). We first noticed that LK cells proliferated faster than LSK cells (Fig. 5C). Addition of IL-6 did not affect LK cell proliferation but promoted LSK cell proliferation. In LK+ sorted cells, only 2.8% of LSK were recovered after 7 days while 13.9% of LSK+ cells were recovered in the presence of IL-6 (Fig. 5D). Conversely, in the LSK+ sorted cells cultured for 7 days, we determined the emergence of 40.7% and 19.7% of LK in absence and presence of IL-6 respectively.

Altogether, these observations suggest that HSCLs are a heterogenous population with distinct sensitivity to cytokine signaling. Particularly, IL-6 signaling seems to promote LSK cells by stimulating their proliferation, modify the balance between LSK/ LK by inhibiting the transition of LSK into LK cell state and inversely promote LK into LSK.

### HSCL cells can differentiate and self-renew *in vitro*

HSCL exhibited a molecular signature resembling adult immature HSPC (Fig. 3 and Fig. S2). Based on CFC assays, HSCL harbored erythroid differentiation, and immature mixed cell lineages colonies (CFU-GEMM) (Fig. 6A), confirmed by morphology in MGG staining (Fig. 6B). CFU-GEMM frequency was estimated in CFC assays with various starting cell dilutions (1×10^3^, 5×10^3^ and 10×10^3^ cells). We estimated the frequency of CFU GEMM at 0.004 (1 cell out of 250) (Fig. 6C). Interestingly, in absence of doxycycline, this frequency was similar (Fig. 6C) even though the total number of colonies was dramatically reduced (data not shown), suggesting that BCR ABL promotes the self-renewal of HSCL by enhancing their cell proliferation rate without altering their clonogenicity and differentiation potential.

**Figure 6.**
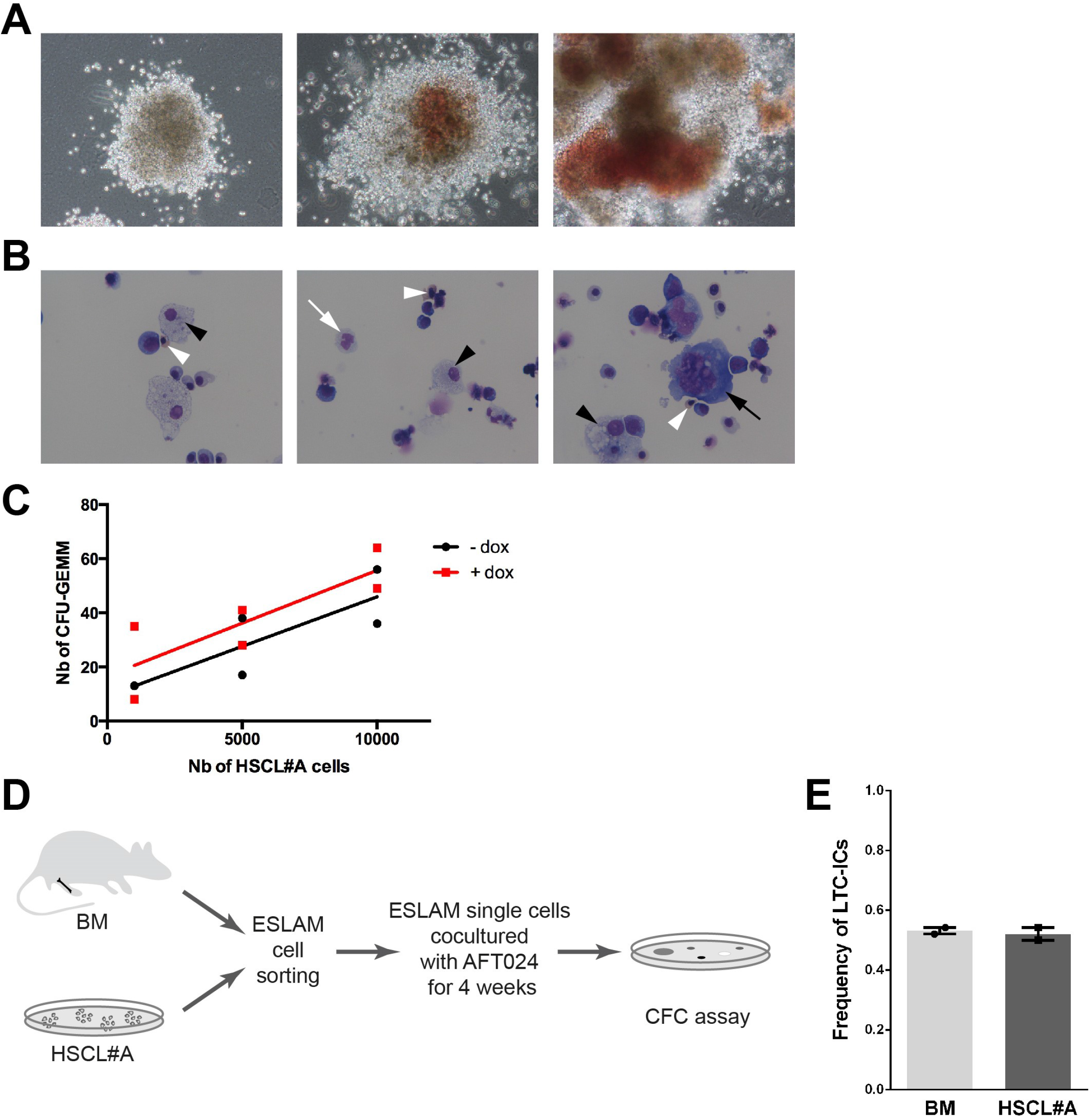
Differentiation potential of HSCL#A cell line. **(A)** Representative colonies obtained from HSCL#A cell line in CFC assay. Distinct morphologies of colonies exhibiting either poor differentiation (upper panel) or differentiation as exemplified with the production of erythroid cells (middle and lower panels). **(B)** Cytospin preparation of a single GEMM colony counterstained with May- Grunwald-Giemsa. Monocytes/macrophages (upper panel), granulocytes (middle panel), megakaryocytes (lower panel) and erythrocytes can be detected. **(C)** Regression analysis of the number of GEMM colonies according to the number of HSCL#A cells. 10^3^, 5×10^3^ and 10^4^ HSCL#A cells were plated in methylcellulose and grown for 2 weeks in duplicates. CFC assays were performed in presence (red, squares) or absence (black, circles) of doxycycline. R-squared values are 0.7224 in absence and 0.6856 in presence of doxycycline. **(D)** Schematic representation of the LTC-IC experimental design. **(E)** Frequency of LTC-ICs isolated from ESLAM bone marrow and HSCL#A cells (n=2 experiments).

Having established that HSCL have erythroid and myeloid potential, we next assessed their B and T lymphoid potential (Fig. S3A, B). HSCL#A cells were cocultured on OP9 or OP9-DL4 stromal cells in presence of IL-7 and Flt-3 cytokines during 14 days. We used fetal liver LSK cells as positive control. We analyzed by flow cytometry the production of pre-B (IgM-B220+CD43-) and pro-B cells (IgM-B220+CD43+), CD4+, CD8+ and CD4/CD8 double-positive T lymphocytes. While fetal liver LSK cells differentiated well into Pro-B cells (on OP-9) and T cells (on OP9-DL4), HSCL#A cells did not differentiate towards lymphoid cell lineages.

The ability of HSCL to self-renew toward long-term hematopoietic potential was measured *in vitro* in LTC-IC (long-term culture-initiating cell) assay. EPCR+CD45+/loCD48-CD150+ (ESLAM) cells were sorted at a single cell level and co-cultured on AFT024 feeder layer during 4 weeks. LTC-IC-derived progenitors were assessed by CFU assay (Fig. 6D). The frequencies of LTC-ICs within ESLAM cells from adult bone marrow and from HSCL#A were respectively 0.53 and 0.52 (1 cell out of 2) (Fig. 6E).

## DISCUSSION

We report here for the first time, an embryonic stem-cell-derived inducible hematopoiesis model under the control of BCR-ABL Tyrosine kinase activity. BCR-ABL, along with HOXB4 ^52^ and STAT5 ^53^, is the only oncogene which has been shown to induce HSC potential in mESCs ^5,6^. However, in all experimental studies reported to date, a retrovirus-mediated enforced expression of BCR-ABL was required and this rendered the precise molecular mechanisms involved in hematopoiesis difficult ^5,6^. There is only one mouse model showing the oncogene-dependence of hematopoiesis in a Sca-1-transgenic mouse ^54^ but no ESCs-derived BCR-ABL-dependent hematopoiesis has been reported so far. In this study, we used a Tet-On inducible system to tightly control the expression of *BCR-ABL^P210^* in an ESC-based model of hematopoiesis. As opposed to previous studies, a single copy of BCR-ABL coding sequence was inserted by homologous recombination into an inducible expression genetic system. The inducible nature of this genetically engineered ESC model allows a tight control of BCR-ABL expression at different stages of hematopoietic differentiation from an undifferentiated ESC state. We observed that the expression of BCR-ABL starting from the undifferentiated ESC stage reduced the number of hematopoietic progenitors but did not affect the distribution of CFC colonies (Fig. 1). This is consistent with previous reports where BCR-ABL retroviral transduction in ESCs altered their differentiation and the formation of hematopoietic precursors by modulating STAT3 activity, a key regulator of ESC self-renewal ^55,56^. On the contrary, the expression of BCR-ABL starting from day 5 of ESC differentiation enable the generation of immature hematopoietic progenitors with *in vitro* multilineage differentiation potential. The increase of hematopoietic colonies consecutive to BCR-ABL expression result from a bias toward hematopoietic differentiation and/or an altered regulation of cell proliferation/survival which is a common role assumed by the oncogene in CML myeloproliferative disease (for reviews, see ^57,58^). Interestingly, the decrease frequency of CFU-MK and the increase of CFU-GM were also observed when BCR-ABL was expressed from day 10 onwards but not when it was expressed between day 5 and day 10. These observations suggest that BCR-ABL affects the differentiation potential of more mature hematopoietic progenitors. Strikingly, the increase of CFU-GEMM colonies was only observed when BCR-ABL was expressed from day 5 onwards but not in the situations where it was expressed between day 5 and day 10 or after day 10. This could reflect a combined role of BCR-ABL in the emergence of immature hematopoietic progenitors and their cell survival. Taken together, our timely controlled inducible system unravels multiple roles of BCR-ABL during ESC differentiation in each wave of hematopoiesis, during the hemangioblast stage (before day 5) and at the stage of primitive and definitive HSC emergence (from day 5 to 10).

It is generally assumed that the constitutive tyrosine kinase activity of BCR-ABL oncoprotein promotes CML cell growth and cell survival leading to relative cytokine independence ^59^. Conversely, several cytokines have been reported to expand primitive CML cells including IL-6 ^60–63^. IL-6 is highly expressed in CML patients ^64^ and its expression is associated with poor prognosis ^65,66^. IL-6 signaling has also been proposed to regulate LSC quiescence through the regulation of JAK1-STAT3 ^67^. Lastly, IL-6 is thought to alter the differentiation of LSCs and normal HSPCs towards myeloid lineage at the expense of lymphoid cell lineage ^61,62^. Here, we generated several hematopoietic cell lines which exhibit a certain heterogeneity between LK and LSK subpopulations. We found that IL-6 cytokine signaling promoted LSK cell proliferation but not LK. Addition of IL-6 also stimulated the reappearance of LSK cells from the LK subpopulation and reciprocally inhibited the emergence of LK cells from the LSK subpopulation. Taken together, our results suggest distinct roles of IL-6 in cell heterogeneity and proliferation of immature hematopoietic cells.

Interestingly, upon maintenance of BCR-ABL expression, we succeeded in isolating hematopoietic cell lines (HSCL) that express markers of immature hematopoietic stem cells but with a distinct transcriptomic profile compared to fetal and adult LSK. We also demonstrated that these cells functionally exhibit *in vitro* selfrenewal and differentiation properties.

The signaling pathways involved and required for induction of HSC potential from ESC or iPSC remain elusive. The model that we report here could be of major interest to uncover these pathways that could then be applied to the strategies of HSC generation from iPSC for transplantation purposes.

Altogether, our system provides a novel useful hematopoietic leukemic stem cell model for drug screening assays in CML and in other hematological malignancies, and highlight the molecular mechanisms and self-renewal pathways in PSC-derived HSCs.

## ACKNOWLEDGMENTS

We thank Konrad Hochedlinger for KH2 ESC and plasmids for gene targeting, Julie Chaumeil for OP9 and OP9-DL4 cells, Warren Pear for the pMIGR-BCR-ABL plasmid. We are very grateful to the transcriptomic platform GENOM’IC at Institut Cochin. This work was supported by the ATIP/AVENIR program, ANR INGESTEM Infrastructure, INSERM and Paris Saclay University. A.Z., S.S. and Y.I. received funding from Vaincre le Cancer NRB. I.S. received funding from the Ministère de l’Enseignement Supérieur de la Recherche et de l’Innovation.

## AUTHORSHIP CONTRIBUTION

J.A., A.F. and A.T. designed the experiments; J.A., A.Z., I.S., D.C., L.E.B.S., S.S., Y.I., S.C. and A.F. performed the experiments; J.A., A.Z., I.S., C.D., A.B.G., A.T. and A.F. analyzed data; J.A., I.S., C.D., A.B.G., A.T. and A.F. wrote the manuscript.

