## supplementary material for "BCR-ABL promotes hematopoietic stem and progenitor cell formation in embryonic stem cells"

**Supplementary Figure 1. Generation of BCR-ABL$^{p210}$ inducible ESC. (A)** Schematic representation of the genetically modified *ROSA26* and *Cola1* loci. **(B)** Immunodetection of p-BCR-ABL, p-CRKL and β-actin by western-blot in ESC cultured in absence or in presence of doxycycline for 24h. 20 µg protein extracts were used together with PathScan® Bcr/Abl Activity Assay antibodies (5300S, Cell Signaling Technology).

**Supplementary Figure 2. HSCL#C cells express surface markers characteristic of immature hematopoietic progenitors. (A)** Representative FACS profiles of CD48-CD150+ fraction in LSK (Lin-CD117+Sca1+) and LK (Lin-Sca1-CD117+) cell populations. **(B)** Representative FACS profile of CD48-CD45+EPCR+ cell populations. The gating strategies were designed according to the bone marrow samples.

**Supplementary Figure 3. Lymphoid differentiation potential of HSCL#A cell line.** Representative FACS profiles using a combination of surface markers to analyze B cell linage (A) and T cell (B) lineage differentiation. **(A)** Upper panels: the gating strategy for FACS analysis was designed to determine B cell progenitors using adult bone marrow samples. We analyzed CD43 expression in the B220-/loIgM- cell population. Middle panels: cell sorted LKS from fetal liver were cultured for 2 weeks on OP9 stromal cells in presence of Il-7 and Flt-3. Lower panels: HSCL#A cells were cultured in the same conditions in presence of doxycycline. **(B)** Upper panels: A similar strategy was used to analyze T cell differentiation based on the analysis of the expression of CD4 and CD8 cell surface markers in the B220- cell population. Middle panels : cell sorted LKS from fetal liver were cultured for 2 weeks on OP9-DL4 stromal cells in presence of Il-7 and Flt-3. Lower panels: HSCL#A cells were cultured in the same conditions in presence of doxycycline.

**Supplementary Table 1. List of antibodies used for flow cytometry.**

| Antibody | Catalog Number | Company |
|---|---|---|
| CD45 APC-Cy7 | 103154 | BioLegend |
| CD150 PE-Cy7 | 115914 | BioLegend |
| CD201 (EPCR) PE | 12-2012-82 | eBioscience |
| CD48 APC | 17-0481-82 | Invitrogen |
| CD34 BV421 | 343610 | BioLegend |
| CD41 FITC | 11-0411-82 | eBioscience |
| TER-119 PE-Cy5 | 15-5921-82 | eBioscience |
| Mac-1 (CD11b) PE-Cy5 | 15-0112-82 | eBioscience |
| Gr-1 PE-Cy5 | 15-5931-82 | Invitrogen |
| B220 (CD45R) PE | 103208 | BioLegend |
| B220 (CD45R) PE-Cy5 | 15-0452-82 | eBioscience |
| B220 (CD45R) PE-Cy7 | 552094 | BD Biosciences |
| CD3e | 15-0021-82 | eBioscience |
| c-Kit (CD117) APC-eFluor780 | 105826 | BioLegend |
| CD43 FITC | 143203 | BioLegend |
| CD4 FITC | 100406 | BioLegend |
| CD8a BV421 | 115910 | BioLegend |
| Fc Block (CD16/32) | 553142 | BD Biosciences |

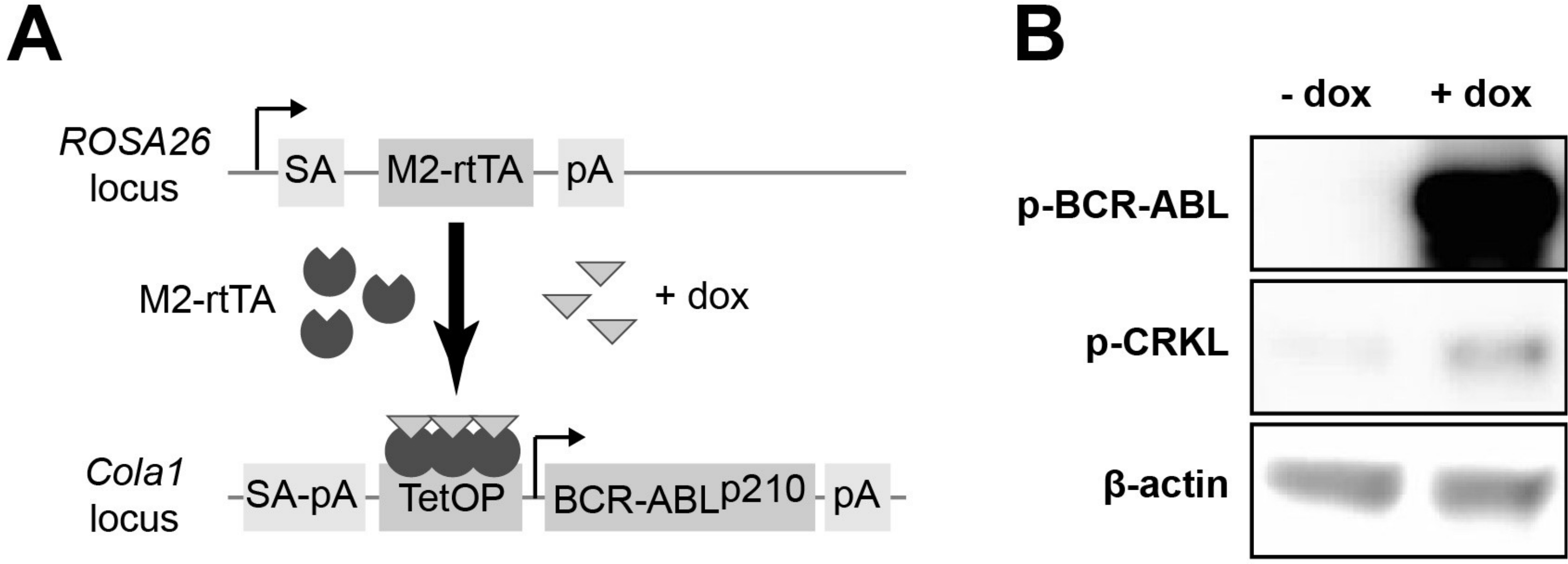


### Figure S1 (Artus et al.)

**A**

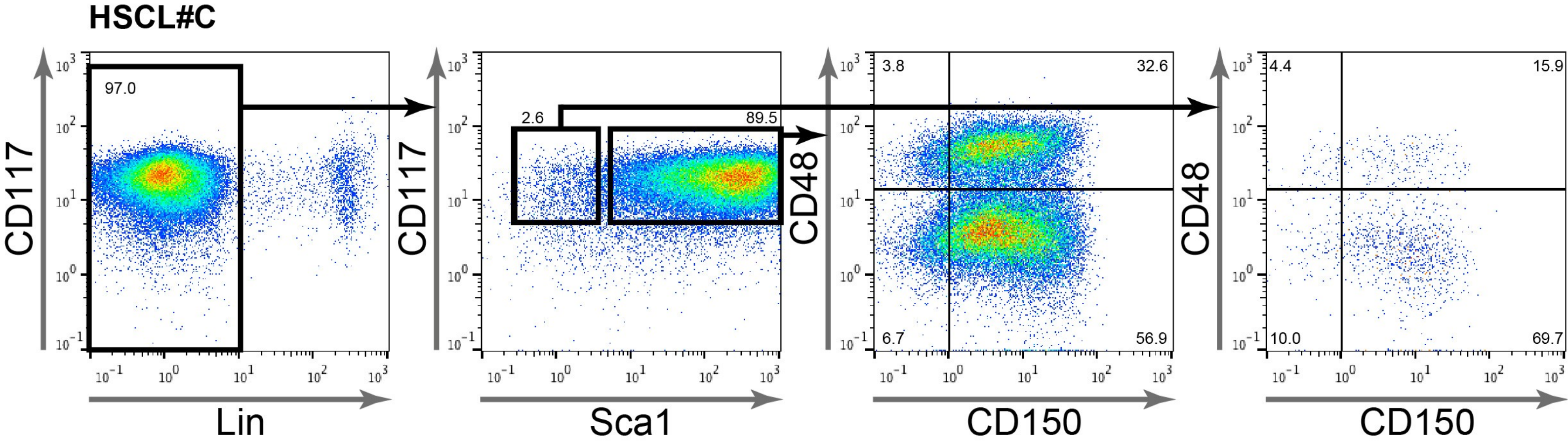


**B**

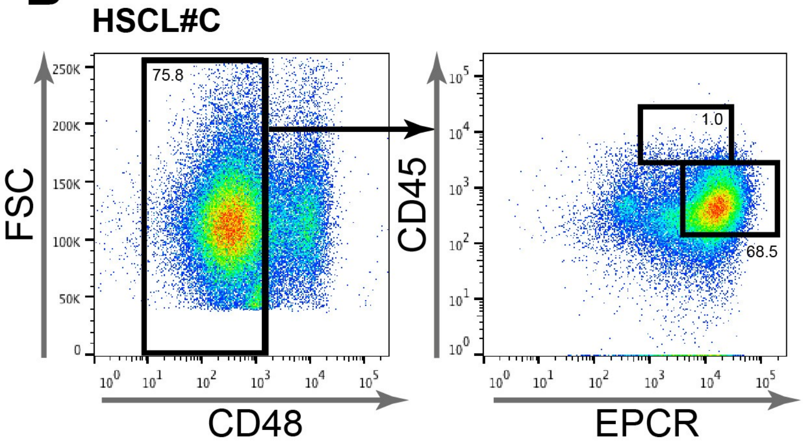


**Figure S2 (Supplement of Fig.3) (Artus et al.)**

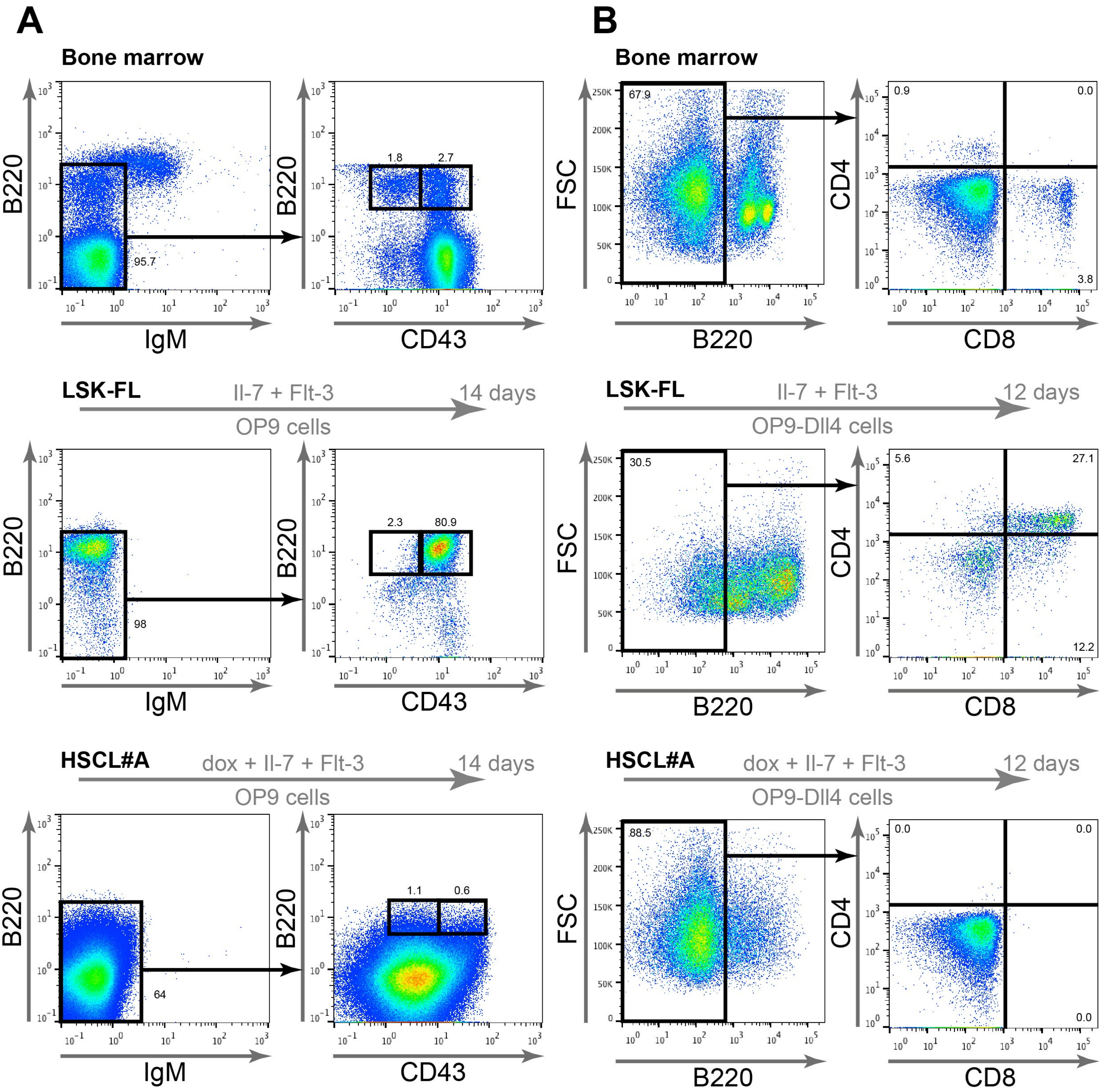


**Figure S3 (Supplement of Fig. 6) (Artus et al.)**
